# Properties of DNA in concentrated aqueous solutions of LiCl suggest transition to C-DNA

**DOI:** 10.1101/2024.09.20.613475

**Authors:** Alexey K. Mazur, Mounir Maaloum, Eugene Gladyshev

**Affiliations:** Université Paris Cité, CNRS, Laboratoire de Biochimie Théorique, Paris, France; Institut Pasteur, Université Paris Cité, Molecular Genetics and Epigenetics Unit, Paris, France; Institut Charles Sadron, CNRS, Université de Strasbourg, Strasbourg, France

**Keywords:** DNA, C-DNA, circular dichroism, atomic force microscopy

## Abstract

C-DNA represents a canonical DNA form related to B-DNA. While C-DNA is known to exist in air-dried fibers, its occurrence in aqueous solutions remains subject to debate. In fibers, the transition from B- to C-DNA is promoted by reduced hydration in the presence of certain monovalent cations (most notably Li+), and this process is generally associated with an increase in the helical twist. To understand if the B-to-C transition can occur in liquid media in principle, we analyzed properties of several circular DNA substrates in aqueous solutions with varying concentrations of LiCl (0.1-8 M). To this end, we estimated changes in the helical twist directly from circular dichroism (CD) spectra of a large supercoiled plasmid and, in parallel, assayed conformational changes in DNA minicircles by atomic force microscopy (AFM). We found that the helical twist increased continuously over the entire range of tested LiCl concentrations, without being subject to saturation even at 8 M LiCl. The overall increase in the helical twist was compatible with the B-to-C transition occurring in solution at a concentration of LiCl of about 8 M, suggesting that C-DNA should be stable above this level.

## INTRODUCTION

C-DNA is a canonical right-handed double-helical form of DNA discovered in the 1950s along with A- and B-DNA by molecular modeling based on X-ray fiber diffraction and other experimental data (1–3). While C-DNA features characteristic diffraction patterns and helical parameters, its putative conformations are not separated by large energy barriers from B-DNA, and, thus, C-DNA has often been considered as a close relative of B-DNA (2, 4–6). Yet, whereas A- and B-DNA can be found *in vitro* and *in vivo*, C-DNA was only convincingly demonstrated in fibers under low air humidity (2, 7), and its existence in aqueous solutions, along with its biological role, remained subject to debate.

Recently, the ability of DNA to undergo the B-to-C transition was proposed to facilitate recombination-independent recognition of DNA homology (8, 9). The existence of such a mechanism was first illuminated by genetic studies of two genome-defense processes in the fungus *Neurospora crassa*, known as repeat-induced point mutation (RIP) and meiotic silencing by unpaired DNA (MSUD) (9, 10). Analysis of RIP and MSUD suggested that DNA homology was detected in a way incompatible with the involvement of single-stranded intermediates, and, instead, relying on the direct pairing of intact double-stranded DNAs (dsDNAs) (9, 10). To account for those genetic observations, an allatom model was proposed, in which homologous dsDNAs could align by a series of major-groove contacts, with each contact composed of a stack of 3-4 planar quartets, and each quartet formed by two identical Watson-Crick base-pairs (11). According to the model, such contacts must be spaced with the periodicity close to the helical repeat, and the overall pairing can be further enhanced by having the participating dsDNAs coiled into a right-handed plectoneme (11). By setting the inter-contact length to the optimal values obtained for RIP and MSUD, it was determined by molecular dynamics simulations that the conformation of the intervening segments should be strongly shifted towards C-DNA (9).

Overall, two properties of C-DNA make it particularly fit for homologous pairing. First, the shallow major groove of C-DNA permits almost perfect initial quartet contacts without atom-atom clashing (8). Second, the helical repeat of approximately 9 bp enables a pair of C-DNA duplexes coiled into a right-handed plectoneme to have their major grooves facing each other every 22 bp, thus allowing the high density of homologous contacts to detect even relatively short repeats (8, 10). Given these considerations, the B-to-C transition emerges as an attractive strategy to lower the energy barrier for the formation of quartet stacks during homologous dsDNA-dsDNA pairing.

Until the 1990s, C-DNA remained a subject of active investigations. Purported B-to-C transitions were routinely monitored by measuring the circular dichroism (CD) of DNA in the near-UV (180-320 nm) range (12–15). The reference CD spectra of B- and C-DNA were obtained from unoriented DNA films under conditions regarded as similar to those used in the original X-ray fiber diffraction studies (2, 16). At high air humidity, those spectra agreed with the results obtained in solution (16). Using this approach, partial B-to-C-DNA transitions were found in various viruses (12, 13), protein complexes (17, 18), chromatin (14), as well as in concentrated aqueous solutions of common salts such as NaCl (19, 20) and in the presence of dehydrating co-solvents (19, 21). However, these CD-based results were challenged by other methods (22–25), and this controversy could not be resolved at that time. Eventually, the view prevailed that the original assignment of the reference CD spectra was incorrect, and that the DNA conformation giving rise to the C-form spectrum corresponded to “a minor variant of the B-form” (22, 23, 26, 27). However, the issue remained unsettled, as similar CD effects, observed in other situations, continued to be interpreted as evidence for partial B-to-C DNA transitions (6, 28, 29).

In this study, we analyzed the CD of supercoiled circular DNA in concentrated aqueous solutions of LiCl to elucidate the effect of LiCl on the helical twist, a critical parameter that differentiates B- and C-DNA. Our approach relied on the two established notions. First, CD provides a sensitive reporter of the transitions between the canonical DNA forms (6, 30), and, at moderate salt concentrations, its near-UV amplitude clearly correlates with the helical twist (31). Second, the CD of supercoiled DNA is known to vary smoothly with the degree of supercoiling (32), thus offering a possibility to probe the helical twist directly (33). Two types of circular DNA were analyzed: a 2877-bp plasmid and a 178-bp minicircle. In low salt, due to their small size, the minicircles were expected to exist as rigid flat rings resisting supercoiling (34), while in high salt they were expected to deform due to excessive twisting. These deformations were detected by AFM, as an orthogonal approach to CD.

We used LiCl for three reasons. First, although Li^+^ is not biologically relevant, it is the only simple cation that promotes the B-to-C transition while attenuating the B-to-A transition in both fibers and solution (25). Second, aqueous LiCl solutions up to 16 M remain stable, the property that allows the water activity to be reduced to 11% (35), which covers the entire range of air humidity used in X-ray fiber diffraction experiments. Third, concentrated LiCl solutions remain amenable to near-UV CD spectroscopy. These properties allowed us to estimate changes in the helical twist directly from CD spectra of supercoiled DNA in solution and under the conditions of low water activity expected to favor the B-to-C transition. By assuming that similar secondary structures produce similar CD spectra of polymer DNA, we were able to evaluate the changes of the helical twist by measuring the salt-dependent difference between CD spectra of two circular topoisomers with different superhelical densities. We found that the helical twist increases steadily for LiCl concentrations between 1 M and 8 M, with the overall increase relative to low-salt conditions reaching 1.6 ± 0.15°. In DNA fibers, this result would correspond to a decrease in the helical pitch from the canonical 10.0 to 9.6 bp/turn. By X-ray fiber diffraction experiments, the latter value was found in both B- and C-DNA conformations corresponding to the onset of the B-to-C transition (36). Taken together, our results suggest that the B-to-C transition should be favored in aqueous solutions with LiCl concentrations above 8 M.

## MATERIALS AND METHODS

### Plasmid construction

Plasmid pEAG270A was produced by inserting the following 212-bp DNA fragment into pUC19 between EcoRI and XbaI (*loxP* sites are underlined; the recognition site of Nt.BsmAI is in bold): <u>ATAACTTCGTATAATGTATGCTATACGAAGTTAT</u>ACCGGTCAATGCCTGCATGGCAATCATATCAAAAGGG CATTGAGCAGGTACCTTTCCTCTCTTAGCTCGGCTAACAGTCAGGACGACAAACCACGCGTAAA**GTCTC**AC GATTGGATCTGCTCATGAAGGTAGGCGATAACTAGT<u>ATAACTTCGTATAATGTATGCTATACGAAGTTAT</u> The inserted fragment was verified by Sanger sequencing. The plasmid was propagated in DH10B cells.

### Purifying negatively supercoiled form of pEAG270A (P_S_)

10 ul of the plasmid stock culture were inoculated into 400 ml of LB medium with ampicillin (100 ug/ml) and grown overnight at 37 °C / 180 RPM. Plasmid DNA was prepared by the alkaline lysis method. Supercoiled DNA (P_S_) was separated on a 0.7% agarose gel containing ethidium bromide, electroeluted, concentrated by anion exchange chromatography using a HiTrap Q HP 1-ml column (Cytiva), precipitated with ethanol, resuspended in water, and quantified using NanoDrop 2000. For the CD analysis, DNA was precipitated with 1 M LiCl and ethanol, and resuspended in water to the final concentration of 1 ug/ul.

### Obtaining topoisomers of P_S_ (pEAG270A)

To obtain closed circular (P_C_), open circular (P_O_), and linear (P_L_) forms of pEAG270A, raw plasmid DNA was treated with Vaccinia topoisomerase I (produced in the lab), Nt.BspQI (NEB), and ScaI (NEB), respectively. All products were purified exactly as P_S_.

### Obtaining closed minicircles (M_C_)

Raw plasmid DNA was treated with Cre recombinase (produced in the lab) and then ScaI (NEB), to liberate excised minicircles. The latter were purified by gel-filtration chromatography on a Superose 6 Increase 10/300 GL column (Cytiva) and concentrated by anion exchange chromatography using a HiTrap Q HP 1-ml column (Cytiva). Preparations of minicircles were subsequently treated with BAL-31 nuclease to remove nicked or single-stranded byproducts. Stopped reactions were loaded directly on a Superose 6 column, and intact minicircles were purified as a single major peak. Mini-circles were precipitated with ethanol, resuspended in water, and quantified using NanoDrop 2000. For the CD analysis, they were precipitated with 1 M LiCl and ethanol, and resuspended in water to the final concentration of 0.5 ug/ul.

### Obtaining nicked minicircles (M_O_)

Intact minicircles (M_C_) were treated with Nt.BsmAI (NEB), purified twice with phenol-chloroform, precipitated with ethanol, resuspended in water, and quantified using NanoDrop 2000. For the CD analysis, nicked minicircles was precipitated with 1 M LiCl and ethanol, and resuspended in water to the final concentration of 0.5 ug/ul.

### Circular Dichroism

All CD experiments were done at the Molecular Biophysics facility (Institut Pasteur) using a J-1500 spectrophotometer (Jasco) equipped with a PM-539 detector. The following parameters were used: range = 400-220 nm, data pitch = 1 nm, CD scale = 200 mdeg/0.1 dOD, FL scale = 200 mdeg/0.1 dOD, D.I.T. = 2 sec, bandwidth = 1.00 nm. A stock of 10 M LiCl was used. Plasmid and minicircle samples were measured under similar conditions with the following differences. Plasmid samples: DNA = 10 ug / 10 ul, total volume = 400 ul, cuvette = 104F-QS (Hellma), accumulations = 6. Mini-circle samples: DNA = 2.5 ug / 5 ul, total volume = 75 ul, cuvette = 105.202-QS (Hellma); accumulations = 12. Blank solutions with matching LiCl concentrations were used to establish baselines for correction. Normalized raw CD data were used for analysis and plotting (no filtering was applied).

### Atomic Force Microscopy: Sample preparation and scanning

DNA was diluted to 2 ug/ml in a buffer containing different concentrations of LiCl. Samples were prepared on freshly cleaved mica discs. Following cleavage, the mica was treated with a solution of NiCl_2_ for 10 seconds, rinsed with ultra-pure water, and dried with a stream of dry nitrogen. A 5-ul droplet of DNA solution was deposited onto nickel-treated mica surface for 1 min. The surface was rinsed with 0.01% uranyl acetate in order to stabilize the DNA molecules for AFM imaging in air. Sample were then rapidly rinsed with pure water (Millipore) to obtain a clean surface after drying with filter paper.

Images were obtained by scanning the samples using a Dimension-Icon AFM microscope (Bruker) operated in peak-force tapping mode. Ultra-sharp silicon cantilevers (ScanAsyst 0.4 N.m-1) with a nominal tip radius < 5 nm were used. AFM images were acquired under in air at room temperature. The force was reduced to avoid dragging molecules by the tip. Integral gain was adjusted to yield sharp images. No online filtering was applied. Images were subsequently processed by flattening to remove the background slope.

## RESULTS AND DISCUSSION

### Supercoiling impedes salt-induced changes in CD

The preparation of DNA constructs and the abbreviations used in this study are explained in Figure 1A-D. Capital letters P and M refer to the 2877-bp plasmid and the 178-bp minicircle, respectively. Minicircles were derived from the plasmid by site-specific recombination. Subscripts S, C, O, and L refer to the negatively supercoiled (native), closed circular, open circular (nicked), and linear forms of P or M, respectively. The near-UV (220-400 nm) CD spectra of DNA were measured in aqueous solutions of LiCl with concentrations ranging from 0.1 M to 8.0 M.

**Figure 1.**
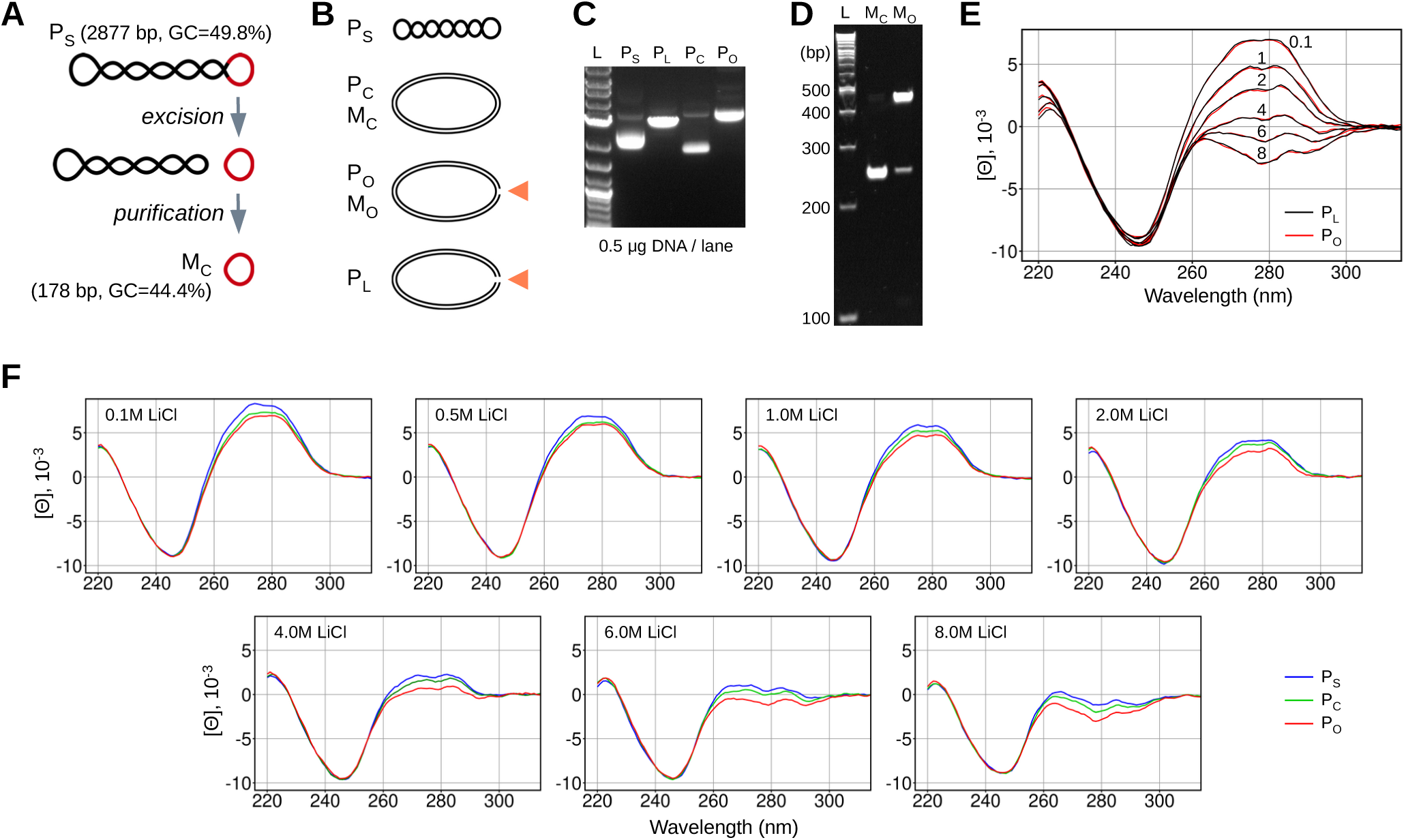
CD spectra of plasmid DNA topoisomers in aqueous LiCl solutions. (**A)** Minicircles were excised from supercoiled plasmid DNA (pEAG270A) with Cre recombinase, treated with BAL-31 to remove nicked or single-stranded byproducts, and purified by gel-filtration chromatography. The DNA contours are not shown to scale. **(B**) Designations of DNA constructs used in the text with sketches of the topology. The contours are not shown to scale. **(C)** Plasmid topoisomers were resolved by agarose gel electrophoresis in the presence of ethidium bromide. P_L_, P_C_, and P_O_ were obtained from supercoiled plasmid DNA (pEAG270A) by treatment with ScaI, Nt.BspQI, and Vaccinia topoisomerase I, respectively. DNA was purified by agarose gel electrophoresis followed by electroelution (Methods). **(D)** Minicircles were resolved by polyacrylamide gel electrophoresis in the presence of chloroquine. M_O_ was obtained from M_C_ by treatment with Nt.BsmAI. **(E)** CD spectra of P_O_ and P_L_ in aqueous LiCl solutions. Concentrations of LiCl are indicated above the plots. **(F)** CD spectra of P_O_, P_C_, and P_S_ in aqueous LiCl solutions. Concentrations of LiCl are indicated. The relative average linking numbers were measured in low salt conditions using the gel band counting method (50) and found to be 0, -0.5 and -16.8 for P_O_, P_C_ and P_S_, respectively.

The CD spectra of P_O_ and P_L_ are shown in Figure 1E. The collapse and inversion of the broad band around 275 nm is a characteristic pattern originally attributed to the B-to-C transition (16, 19, 37). The accompanying changes at shorter wavelengths are not monotonous and probably reflect other factors. At all tested LiCl concentrations, the spectra of P_O_and P_L_ can be superimposed within the errors (38).

Figure 1F compares the same CD spectra of P_O_ with those of P_C_ and P_S_. In 0.1 M LiCl, all of them have typical B-DNA shapes with varying positive intensities of the 275-nm band, ranked as P _O_ < P_C_ < P_S_. Because these constructs represent topological isomers of the same circular DNA, this ranking reflects conformational changes in DNA caused by supercoiling stress, which is proportional to the superhelical density (32). This long-known effect (38) remains poorly understood. While increasing concentration of LiCl appears to affect all three topoisomers similarly, P_C_ and P_S_ feature somewhat smaller changes in the amplitude of the 275-nm CD band (Fig. 1F). As a result, in 8.0 M LiCl, the CD spectra of P_O_, P_C_, and P_S_ are arranged in the same order as in 0.1M LiCl, but with their 275-nm bands becoming inverted and further separated. In this context, the separation between P_C_ and P_O_ increases more than the one between P_C_ and P_S_.

The CD spectra of the 178-bp minicircles (M_C_ and M_O_) were measured in similar conditions and are shown in Figure 2A. In this series, P_L_ was used as a reference, because linearizing minicircle DNA for this purpose was impractical. In 0.1M LiCl, the spectra of M_C_, M_O_, and P_L_ are very similar (Fig. 2A). With the helical repeat of ∼10.5 bp/turn, 178 bp correspond to exactly 17 turns. Therefore, M_C_ is not torsionally strained, and the difference between M_C_ and M_O_ should be negligible. While DNA in minicircles is constantly bent along its entire length, this feature apparently does not affect its CD in the tested range (220-400 nm).

**Figure 2.**
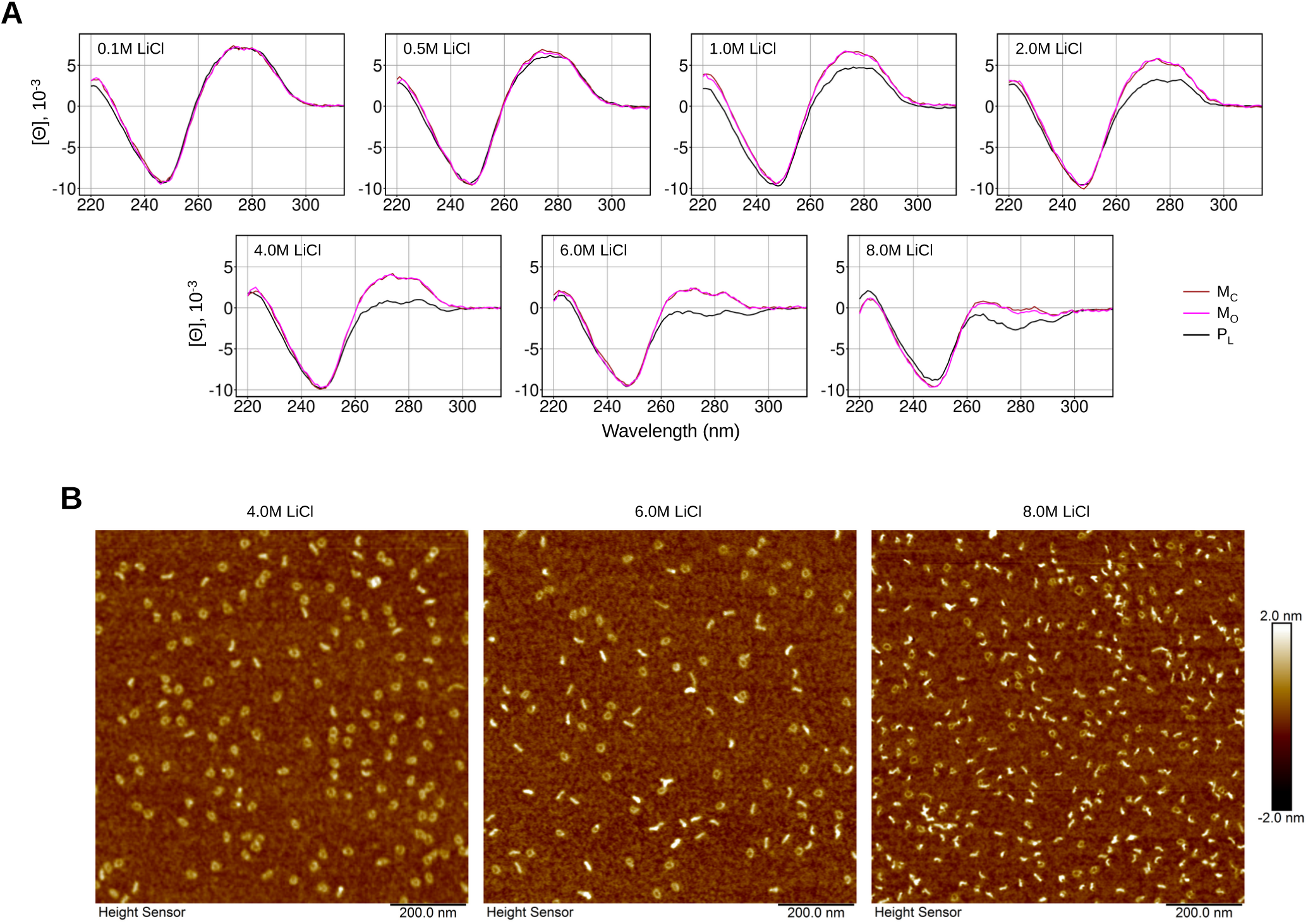
Properties of minicircles in aqueous LiCl solutions. **(A)** CD spectra of closed (M_C_) and nicked (M_O_) minicircles in aqueous LiCl solutions, as compared with the corresponding spectra of P_L_. Concentrations of LiCl are indicated. **(B)** AFM scans of ensembles of minicircles (M_C_) in concentrated LiCl solutions. The corresponding LiCl concentrations are indicated above the images.

Under low salt conditions, closed minicircles (M_C_) are restrained by strong interhelical electrostatic repulsion and represent flat rigid rings that cannot twist or bend (34). In such conditions (below 0.1 M LiCl) the CD spectrum of M_C_ remains constant (Fig. S2), but it starts to change visibly between 0.1 M and 0.5 M LiCl (Fig. 2A). In contrast, DNA of M_O_ can twist even in rigid flat rings, and the spectrum of M_O_ was expected to follow the spectra of P_L_ and P_O_ (Fig. 1E). Remarkably, the opposite result is observed (Fig. 2A). Specifically, the spectrum of M_O_ diverges from the spectrum of P_L_ (and P_O_) and, instead, follows the spectrum of M_C_. In fact, there are hardly any significant differences in the spectra of M_O_ and M_C_ in 0.1-6.0 M LiCl; and only in 8.0 M LiCl does the spectrum of M_O_ start to deviate towards that of P_L_. Thus, while a single-strand break in a large closed circular DNA can remove all supercoiling constraints and transform its CD spectrum into that of linear DNA (Fig. 1E; ref. (38)), it fails to do so in the context of the 178-bp minicircle (Fig. 2A). Notably, while the effect of LiCl on M_C_ is qualitatively similar to that for P_C_, it appears weaker, and the positive intensity of the 275-nm CD band in the M_C_ spectra does not become inverted even in 8.0 M LiCl.

Minicircles have also been particularly useful for understanding the effects of LiCl on the properties of DNA because of their utility for AFM. In low salt conditions, unlike large plasmids, minicircles are conformationally homogeneous. However, as the concentration of LiCl starts to increase, their conformations are expected to depart from the ideal shape, a process that can be visualized by AFM. It was previously shown that nickel-treated mica could readily adsorb DNA from monovalent salt solutions without the addition of other ions (39, 40). We found that this approach could be adapted for DNA in LiCl solutions with concentrations above 2 M, thus allowing direct qualitative analysis of minicircle conformations at high LiCl concentrations.

Figure 2B shows that in 4.0 M LiCl, 75% of minicircles are still ring-shaped (observed as roughly circular spots with a central hole). Another characteristic highly populated structure corresponds to an elongated straight conformation with the length and thickness suggesting a single minicircle collapsed into a straight rod along its diameter. In 6.0 M LiCl, the proportion of rings is reduced to 49% and collapsed structures already begin to predominate. In 8.0 M LiCl, the percentage of rods appears to have increased further, although simple population count is no longer possible due to the presence aggregates of different sizes. Each of the AFM scans (Fig. 2B) also contains amorphous structures without holes which may correspond to tilted deformed rings. Since the concentration of LiCl in the deposited solution was the only varied parameter, we can reasonably assume that these scans reflect differences between solution ensembles of DNA conformations. Repeated scans of the same samples a few days later yielded similar mixtures of structures; but with the higher proportion of aggregates in 8.0 M LiCl. No changes in the CD spectra were detected as well. Consequently, we favor an idea that transitions between the ring and the rod-shaped structures are reversible, and their mixture is in equilibrium. Rod-shaped structures were also seen in 2.0 M LiCl in both M_C_ and M_O_ samples, but the AFM images were much less clear.

Overall, our results indicate that qualitatively similar conformational changes are induced in closed circular and linear DNA upon increasing LiCl concentration (Fig. 1, 2). These changes are markedly impeded by the ring closure constraints and negative supercoiling, pointing to the DNA twisting as the principal affected parameter.

### Structural interpretation of CD spectra

The characteristic CD spectrum of DNA in the near-UV range is produced by the electrons in non-chiral nucleobases under the influence of two external conditions: (1) the chemical chirality of the proximal deoxyribose rings and (2) the overall helicity of the DNA structure (41). The first factor is considered negligible due to the relatively large distances to the chiral sugar atoms, and a very large number of closer atom-atom contacts in the base-pair stacks. Indeed, reasonably good qualitative and sometimes quantitative agreement with experiment is achieved by theoretical predictions of CD spectra that take into account only base-base interactions (42). The helical chirality emerges due to the base-pair twisting, which is the key element of this theory (43). It is logical, therefore, that the influence of supercoiling and solution conditions on CD spectra was initially attributed to altered twist angles (19, 38). However, as new data continued to accumulate, this interpretation became increasingly questionable. It was later argued that, due to a comparatively low intensity of the 275-nm CD band, it may involve a non-negligible contribution from interactions with proximal sugars, and therefore, that it cannot be used as a quantitative reporter of DNA twist and/or an evidence of the C-form (22, 44). We agree with this physical argument, but not with the conclusion drawn.

We assume that the intensity of the 275-nm band in the CD spectrum of DNA is affected, directly or indirectly, by multiple factors; in particular, by the transitions between the backbone conformers BI and BII, which invert the orientation of the phosphate groups and modulate the helical electrostatic field at the stacked bases (5). Due to the relatively low intensity of the 275-nm band, these factors may have important roles, and they still cannot be accounted in *ab initio* calculations of CD spectra (45). However, there are good reasons to believe that all of these factors, in the first approximation, change in concert and in proportion to the twist angle. For supercoiling, this idea follows from the earlier established experimental dependence of CD spectra of circular DNA on the magnitude and sign of the superhelical density. Indeed, negative and positive supercoiling are known to change the intensity of the 275-nm in opposite directions, with the magnitude that varies linearly over a wide range around the zero superhelical density (32). Below we argue that the twist is also involved in LiCl-induced structural changes that affect the CD spectrum of DNA.

Closed loops of long double-stranded DNA are described by a simple topological model with three parameters, namely, linking number (*Lk*), twist (*Tw*) and writhe (*Wr*) (46, 47), which are related by the equation

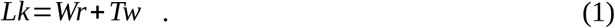

*Lk* is an integer defined as the number of times the two strands are intertwined. It is a topological invariant changed by integer increments, when one of the two strands is broken and rejoined after rotation around the opposite strand. In contrast, *Tw* and *Wr* are real numbers that change with the shape of the loop; they are mutually balanced and are in thermodynamic equilibrium. Mixtures of plasmid topoisomers, such as samples P_C_ and P_S_ produced by topoisomerases or isolated from cells, are characterized by averages ⟨ *Lk* ⟩, ⟨*Wr* ⟩, and ⟨ *Tw* ⟩, all of which are real numbers also related by Eq. (1). Since we will only operate with such averages, the angular brackets will be omitted.

A closed relaxed DNA circle has *Wr* =0 and *Lk*=*Tw_O_* =*Lk_O_*, where *Tw_O_* is the twist in linear DNA or open DNA circles. Note that, according to this definition, *Lk_O_* is a real number that changes continuously. In the harmonic approximation, the minimum energy conformation of open circles is a planar ring (*Wr* =0) which is free from twisting strain (*Tw*=*Tw_O_*) and has the minimum bending strain required for loop closure.

According to Eq. (1), with *Lk* < *Lk_O_*, DNA in flat circles is unwound and torsionally stressed (*Wr*=0 *, Tw*< *Tw_O_*), while without twisting (*Tw*=*Tw_O_, Wr* < 0) it becomes supercoiled, *i.e.*, additionally bent. The supercoiling stress can be relieved by balancing the bending and the twisting strains to obtain a global energy minimum with *Wr* <0 and *Tw* <*Tw_O_*. In general, the excess or deficit in the linking number is partitioned between writhe and a change in twist (from that in linear

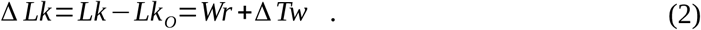

This partitioning was studied with the analysis of electron microscopy (EM) images of supercoiled plasmids and also by Monte-Carlo (MC) simulations. According to the EM data, the proportions of the linking number deficit compensated by writhing and twisting are nearly constant, being *qWr* =*Wr* / Δ *Lk*≈0.72 and *qTw*=1−*qWr* =0.28, respectively, for the natural range of superhelical density *σ*=Δ *Lk* / *Lk_O_* (48). The MC simulations predicted an increase of *qWr* from 0.72 at *σ*≈0 to 0.8 at *σ*≈0.06 (49). Accordingly, the twist of supercoiled topoisomers is

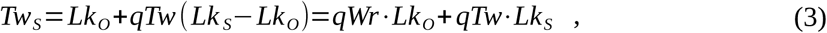

and the twist difference between topoisomers P_C_ and P_S_ is:

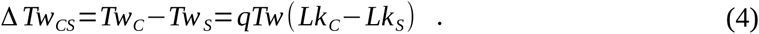

In the above equations, all variables are measured in helical turns. Twisting can also be measured by the average angle between adjacent base pairs, that is, *δ Tw*=Δ *Tw*⋅360 *°* / *N* where *N* is the length of the DNA double helix in base pairs.

Eq. (3) indicates that, when the twist in linear DNA is increased due to the salt effect, it is also increased in supercoiled DNA, but to a lesser extent because of the increased superhelical density. In the linear approximation, these shifts are similar in all of topoisomers and according to Eq. (4) the twist difference between P_C_ and P_S_ should remain constant. Each time the P_S_ twist is increased by *δ Tw_CS_*, it reaches the P_C_ twist value at the previous LiCl concentration. This does not mean that the supercoiled topoisomer P_S_ unfolds and becomes overall similar to P_C_. However, if the CD spectrum of DNA is determined by the helical twist, the corresponding spectrum of P_S_ should be similar to the spectrum of P_C_ in 0.1 M LiCl. This reasoning can be applied to other concentrations of LiCl and topoisomer pairs (P_S_-P_O_, P_C_-P_O_, etc). Overall, the theory predicts that, as the concentration of LiCl increases, CD spectra of all negatively supercoiled topoisomers of the same plasmid DNA should follow the same trajectory, trailing behind the spectra of P_O_ and P_L_.

Figure 3A confirms this prediction. In this example, the spectrum of P_C_ in 0.1 M LiCl is bound by the spectra of P_S_ in 0.1 and 0.5 M LiCl (left panel). Assuming that the CD spectra change smoothly, their intermediate shapes can be predicted approximately as the weighted averages of the boundary spectra. It is then possible to find a concentration of LiCl, at which the spectrum of P_S_matches the target spectrum of P_C_, and then refine it by measuring the CD spectra of P_S_ at and near the predicted concentration (middle panel). Indeed, the P_C_ spectrum at 0.1 M LiCl overlaps within the errors with both the predicted and the actual P_S_ spectrum taken at the predicted LiCl concentration of 0.375 M (middle panel). This result means that, in negatively supercoiled circular DNA, with respect to CD, a reduced linking number can be compensated by an increase in LiCl concentration, and *vice versa*. The linking number, the twist, and the CD spectrum are associated and change concertedly. The CD signal can then be used as a sensitive reporter of these changes, even if the underlying mechanisms relating CD with the structure remain unclear.

**Figure 3.**
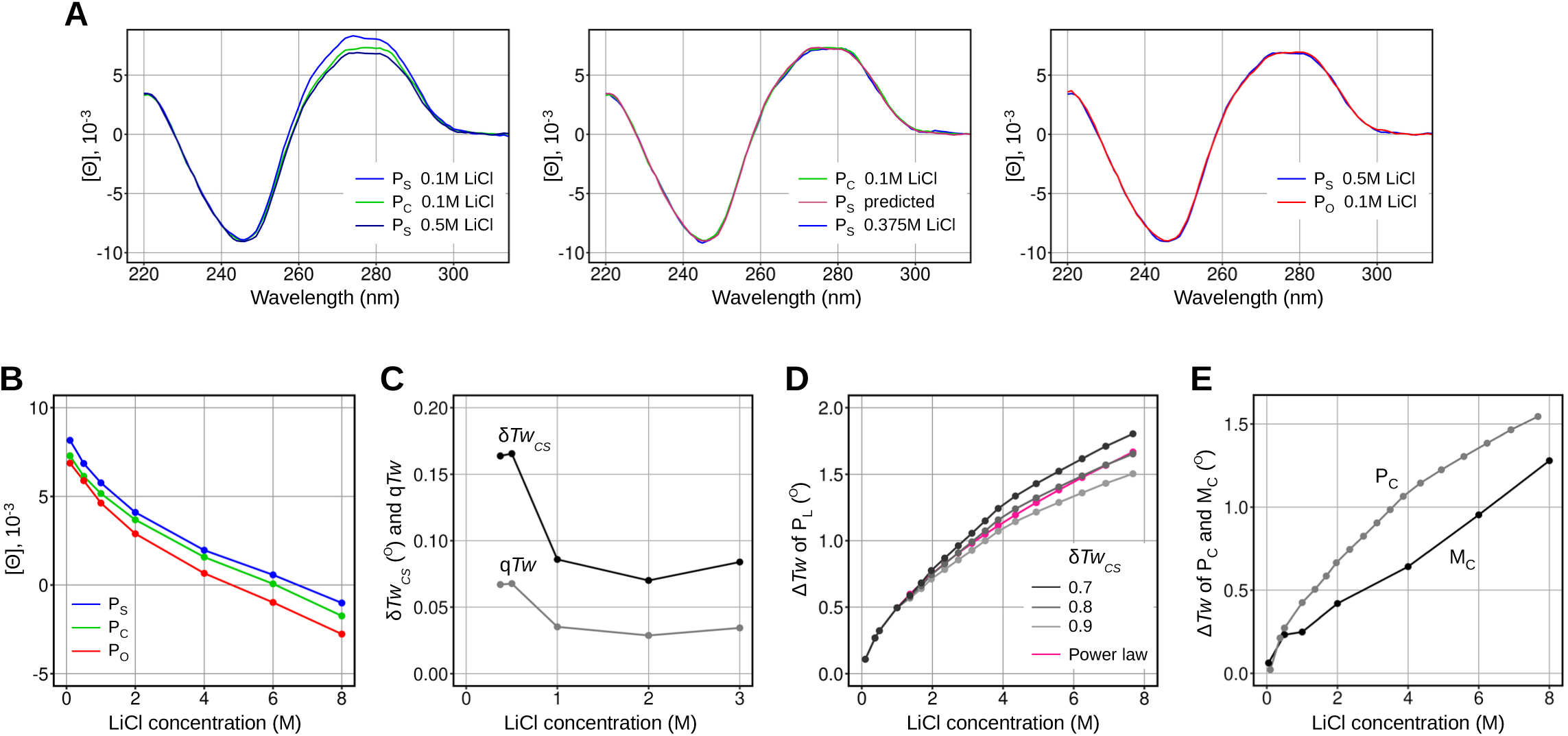
Estimating the differences in the helical twist directly from CD spectra. **(A)** (Left) CD spectra of P_S_ and P_C_ at three LiCl concentrations. The two boundary P_S_ spectra were used to predict the LiCl concentration (0.375 M) at which the spectrum of P_S_ matches the spectrum of P_C_ in 0.1 M LiCl. (Middle) CD spectra of P_S_ and P_C_ in 0.375 M and 0.1 M LiCl, respectively. The interpolated spectrum of P_S_ is also shown. (Right) CD spectra of P_S_ and P_O_ in 0.5 M and 0.1 M LiCl, respectively. **(B)** Dependence of the average CD intensity at 275-278 nm of P_O_, P_C_, and P_S_ on LiCl concentration. **(C)** Dependence of the twist difference between P_S_ and P_C_ on LiCl concentration, as estimated from the comparison of CD spectra. Also shown is the corresponding proportion of the linking number difference compensated by twisting (*qTw*). **(D)** Changes in twist estimated from CD spectra at high LiCl concentrations for the three values of the twist difference between P_S_ and P_C_. The red curve shows the extrapolation of a power law fit to experimental values over the concentration range from 50 mM to 1 M (52). **(E)** Changes in twist in minicircles estimated from CD spectra at high LiCl concentrations, as compared with the similarly measured dependence for P_C_.

According to Figure 3A, the increase in twist upon the transfer of P_S_ from 0.1 M to 0.375 M LiCl is equal to the twist difference between P_S_ and P_C_ in 0.1 M LiCl. This statement is based on a single assumption, namely, that similar CD spectra of this particular circular DNA correspond to similar conformations around the π-electrons of stacked bases. The right panel of Figure 3A shows that the same properties are also inherent in the pair of topoisomers P_S_ and P_O_. Here, the CD spectrum of P_S_ in 0.5 M LiCl and that of P_O_ in 0.1 M LiCl are nearly identical. This fortunate coincidence proves that the CD spectra of P_S_ and P_O\_ behave similarly to the P_S_ and P_C_ spectra in Figure 1F, and that for each P_S_ spectrum one can find an equivalent P_O_ spectrum at a smaller LiCl concentration. The twist of DNA in P_O_ is similar to that of P_L_ (Fig. 1E), therefore, one can use the literature twist value for linear DNA in 0.1 M LiCl as an estimate for P_S_ in 0.5 M LiCl and further refine this concentration if a better estimate is needed. This approach is generally applicable to topoisomers of the same DNA and we use it below to roughly estimate the effect of LiCl on DNA twisting over the entire range of concentrations.

Figure 3B compares the effect of LiCl on the intensity of the 275-nm CD band for P_S_, P_C_, and P_O_ (over an interval covering the positions of the band maximum in all spectra). In low salt, the CD intensities of P_S_ and P_C_ are already well-separated. Their separation decreases at LiCl concentrations up to 1 M, passes through a broad minimum, and then increases again. On the other hand, the CD intensity of P_O_ decreases faster and separates further from P_S_ and P_C_ in higher LiCl concentrations. This trend is qualitatively consistent with the prediction of Eqs. (3) and (4) and with the relationship between changes in the CD intensity and the superhelical density. The separation between P_S_ and P_C_ varies with increasing LiCl concentration, suggesting that these topoisomers may undergo different structural changes that influence the twist/writhe balance.

Figure 3B offers a way to roughly estimate the magnitude of the increase in the helical twist due to increasing the LiCl concentration across the entire range. Based on Eq. (4) and using the approach shown in Figure 3A, we can continue to move towards higher LiCl concentrations by alternating vertical and horizontal steps between the P_S_ and P_C_ plots in Figure 3B. Horizontal steps correspond to finding the next LiCl concentration at which the P_S_ spectrum best matches the P_C_ spectrum at the previous LiCl concentration. Vertical steps correspond to measuring the next target spectrum of P_C_. This algorithm will produce a descending stepwise line plot, with the vertices found on the actual P_S_ and P_C_ plots and every step corresponding to a twist increment of Δ *Tw_CS_*. Using high-resolution graphs (Fig. 3B), the increase in twist could be deduced simply with a pencil and a ruler. Although we have only piece-wise linear approximations of the true dependencies, this procedure still gives a reasonable estimate, as the overall amplitude depends only on the total number of steps.

In principle, the value of Δ *Tw_CS_* can be obtained directly from Eq. (4) using the known value of *qTw* and the linking number difference *Lk_C_*-*Lk_S_* measured independently by the gel band counting method (50). The linking number of open circles is *Lk_O_*=*N*/10.5=274, where N is the plasmid length (2877 bp). In the gel buffer (40 mM Tris-acetate), the corresponding averages for topoisomer distributions of P_S_ and P_C_ were shifted from *Lk_O_* by *ΔLk_C_ =* -0.5 ±0.2 and *ΔLk_S_ =* -16.8 ± 0.3, respectively. Since the LiCl concentration does not affect *Lk_C_* and *Lk_S_*, Eq. (4) gives Δ *Tw_CS_*≈4.6 and the corresponding angular increment per base-pair *δ Tw_CS_*=Δ *Tw_CS_*⋅360/ *n*=0.57 *°*. As shown in the previous section, Δ *Tw_CS_* is equal to Δ *Tw_S_* due to the increase of the LiCl concentration from 0.1 M to 0.375 M. The corresponding increase for linear DNA is found using Eq. (3) as

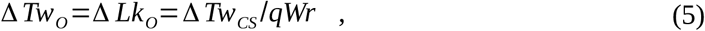

which gives approximately 6.4 of helical turns and *δ Tw_O_*=0.80*°* in terms of base-pair twisting. This last value cannot be correct, as it is at least five times higher than previous consistent estimates obtained by different methods (51, 52). This discrepancy has two plausible, non-mutually exclusive explanations. First, in our experiments, the qTw value may be somewhat lower than the previously estimated range of 0.20-0.28, due to specific salt conditions, and because the coarse-grained models do not account for the distinction between twisting and writhing for short DNA segments. This issue is further discussed in the Supplementary Information section. Second, P_S_ may also collapse into a tightly interwound plectoneme (34, 53–55). For native superhelical densities, such transitions occur at ionic strengths below 1 M (55) and should be accompanied by a strong increase of the negative writhe and a corresponding increase in twist (34). Indeed, the separation between the spectra of P_S_ and P_C_ diminishes in the 0.1-1.0 M range of LiCl concentrations (Fig. 3B). Evidently, in this range, the CD signal decreases faster for P_S_, thus corresponding to a faster increase in twist.

For our present purposes the above uncertainty is not critical. At concentrations above 1.0 M, the P_S_and P_C_ plots become more parallel, and further changes in twist can be approximated by continuing to treat *δ Tw_CS_* as a constant and using a different, more reasonable estimate for it, which can be obtained from the literature data for linear DNA. To this end, we predicted for all P_C_ and P_S_ spectra the LiCl concentrations of good matches with P_O_ and then estimated the corresponding values of *δ Tw_CS_* with a literature three-parameter power-law expression fit to direct experimental values (52). The resulting plot in Figure 3C reveals a significant decrease of *δ Tw_CS_* between 0.5 M and 1.0 M LiCl with a qualitative correlation with fluctuations of the vertical shift between the spectra of P_C_ and P_S_ in Figure 3B. Although the data points at 2.0 M and 3.0 M in Figure 3C correspond to extrapolations beyond the range of concentrations studied experimentally (52), we assume that from 1.0 M to 8.0 M LiCl the *δ Tw_CS_* value is found between 0.07° and 0.09°, which corresponds to *qTw*≈0.04.

Figure 3D shows predictions of the helical twist at high LiCl concentrations based on our CD data. As explained above, a stepwise line plot is created to relate the values P_C_ and P_S_ (Fig. 3B) and scaled with *δ Tw_CS_* values of 0.07°, 0.08° and 0.09°, respectively. This procedure yielded three plots of increasing twist with LiCl concentration, which were then shifted vertically to match the experimental value at 1 M (52). In Figure 3D, these dependencies are compared with the power law (52) extrapolated beyond 1 M. All these plots smoothly continue the experimental data for lower concentrations, and with *δ Tw_CS_* =0.8 *°* the CD-based graph is surprisingly close to the power-law extrapolation. This agreement indicates that the intensity of the 275-nm band in CD spectra of DNA changes with LiCl concentration in qualitative agreement with the coarse-grained model of supercoiled DNA and the assumed relation with the helical twist. The predicted increase of twist is around 1.1° for the concentration range from 1 M to 7.5 M and 1.6 ± 0.15° for the total increase relative to the reference conditions in 50 mM KCl (52). Although this analysis provides only rough estimates, the concerted behavior of CD spectra of supercoiled topoisomers suggests that the helical twist of DNA increases steadily with LiCl concentrations over 0.1-8 M without reaching a plateau.

### Twist changes in minicircles

Given the conclusions of the previous section, a consistent interpretation of the minicircle spectra in Figure 2A is that the helical twist in minicircles also increases, but less so than in large circles and linear DNA. With some caveats, the approach of the previous section can also be used to roughly quantify this increase. Using linear interpolation between P_O_ spectra for consecutive concentrations of LiCl, we predicted for all M_C_ spectra the closest P_O_ spectra and the LiCl concentrations at which they would be obtained. From these spectra, we then estimated the corresponding twist values using the empirical power law formula for linear DNA (52). The quality of the fit is shown in Figure S2, which compares the best-fit computed P_O_ spectra with the corresponding M_C_ spectra. In 0.5 M and 1.0 M LiCl, the agreement between them is as good as for 0.1 M LiCl in Figure 2A, suggesting that between 0.1 M and 1.0 M, the spectra of minicircles follow the same path as the P_O_ spectra. A small discrepancy in spectrum shapes appears in 2.0 M LiCl and continues to increase gradually at higher salt concentrations (Fig. S1). This pattern indicates that some qualitative changes begin to develop in minicircle structures at LiCl concentrations above 1 M, which can be interpreted using the AFM data (Fig. 2B). At LiCl concentrations below 1 M, minicircle DNA appears as a regular double helix that is indistinguishable by CD from linear DNA at some other LiCl concentration. Yet in collapsed structures, DNA may become deformed near the terminal loops (which have significant statistical weight in minicircles), and also at sites of close DNA-DNA contacts. The discrepancy in spectrum shapes at LiCl concentrations above 1 M (Fig. S1) may reflect an increased proportion of collapsed structures (Fig. 2B).

To measure the writhe, we also need the point of zero linking number deficit (*ΔLk*), that is, the LiCl concentration at which the helical twist in minicircles and linear DNA is the same. For our data, the best approximation of this point is 0.1 M LiCl, at which the corresponding CD spectra were almost identical (Fig. 2A). In 0.01 M LiCl, the spectra of M_C_ and P_L_ diverged in the opposite direction (Fig. 2A, Fig. S2), indicating that the twist was lower in linear DNA compared to minicircles, so the *ΔLk* value changed the sign and became positive. The CD spectra of M_C_ did not differ between 0.01 M and 0.1 M LiCl (Fig. S2), that is, the twist did not change, the writhe remained close to zero at both concentrations, and the emerged positive *ΔLk* became compensated by the positive twist difference from linear DNA, which is consistent with previous findings (34).

The estimated twist of minicircles depends on LiCl concentration as shown in Figure 3E. Between 0.1 M and 0.5 M, it increases similarly to the one of P_C_. The increase almost stops between 0.5 and 1.0 M, and then resumes again above 1.0 M, accompanied by the discrepancy in the shapes of CD spectra (Fig. 2A) and characterized by a growing rate, suggesting that it is driven by a mechanism different from that in P_C_ and linear DNA. According to Eq. (2), the corresponding negative writhe equals the twist increase relative to the zero *ΔLk* state. Measured in helical turns, its value is about 0.07 in 2.0 M LiCl and about 0.6 in 8.0 M LiCl. These observations are also consistent with the AFM data. As already mentioned, the collapse of minicircles is likely reversible. It probably begins at high salt concentrations, when the electrostatic repulsion decreases and allows juxtaposed DNA segments to approach each other. At small distances, ion bridges begin to form, which inverts the repulsion and prompts a transition to a tightly interwound plectoneme. At salt concentration above 1.0 M, such fluctuations can apparently occur with small negative writhe, but the transition should result in a stepwise increase accompanied by a compensatory increase in twist (34). The twist plot in Figure 3E reflects the change in the proportions of rings and rods as well as the direct salt effect.

Quasi-identity of the CD spectra of M_C_ and M_O_ (Fig. 2A) indicates that the stacking energy is high and conformations with small twist changes due to unstacked nicks are rare in M_O_. In large circles with *Wr*≈-0.5 (transitions to conformations with Δ *Lk*=+1), increased twist and positive writhe would be possible, but in minicircles they are prevented by ionic bridges, which must be broken on the way through conformations with zero writhe. The behavior of supercoiled minicircles described here differs significantly from the previous detailed CryoEM study (34), most likely due to much a higher ionic strength and transitions to the tightly interwound plectonemes.

### Contrasting views on the structure of DNA in concentrated ionic solutions

Previous objections to the CD-based evidence for the substantial changes in the structure of DNA in concentrated solutions of common salts (including the formation of C-DNA) followed from results of several studies (22, 23, 36, 51). The most important objection emerged from the X-ray diffraction experiments with swollen DNA fibers (36). In this method, a DNA fiber is immersed in a solvent and allowed to swell in a narrow capillary with sealed ends. Comparing these conditions with those found in a bulk solution requires several assumptions; and it appears that one of such assumptions had previously escaped attention. Specifically, in studies using swollen fibers, ions were assumed to only affect DNA directly, without considering their effects on water activity (*a*_w_). In general, water spontaneously diffuses into areas with lower *a*_w_; thus, *a*_w_ in the fiber must increase during swelling. Therefore, transitions from B-DNA to other forms (favored at lower *a*_w_) should be impeded during or after swelling, as the latter can only preserve the original structure of B-DNA or promote this conformation, as observed in all such experiments without exception (36, 56, 57). Surprisingly, the swelling and the associated B-DNA diffraction patterns were also observed for fibers in 6.2 M and 8.0 M LiCl, corresponding to *a*_w_ of 0.60 and 0.43, respectively (35). This result contradicted the X-ray data for dried LiDNA fibers and, therefore, questioned the reality of salt effects on DNA twist in solution. However, this contradiction may have an alternative explanation, provided below.

Fiber wetting in multi-component solvents (such as solutions of salts) is more complex than in pure water. This process may involve substantial diffusion flows of ions and water, resulting in marked changes in solvent composition. For example, for DNA fibers prepared from a 1 mM salt solution and swelled in pure water (57), the final salt concentration increased to 0.1 M (56). In experiments using concentrated LiCl solutions, DNA fibers likely absorbed ions and could noticeably reduce the amount of LiCl in the surrounding medium. The magnitude of this effect can be estimated from the published data (56, 57). There, the initial volume of the solvent was at least 20 times lager than that of the fiber, but as the fiber volume increased substantially during swelling (57), the final difference likely became several times smaller. Overall, because the ion capacity of the fibers and the possible boundary effects of the capillary on all chemical potentials could not be known with certainty, the results of fiber-wetting experiments should be compared to those done in solution only qualitatively. To control *a*_w_ near DNA, the sample had to be equilibrated with the bulk solution, which could not be ensured experimentally.

The same X-ray approach was used to evaluate another instance of C-DNA formation first seen by CD (23). Specifically, it was found that crosslinking of exocyclic amino groups of bases to various simple amines using formaldehyde altered the CD spectra above 260 nm similarly to concentrated salts (58). Below 260 nm, however, the CD spectra of modified DNA diverged, and this difference increased with the number of modifications. The X-ray analysis showed that, regardless of the CD spectrum, modified DNA assumed a B-form conformation with a nearly canonical twist value (23). These results do not contradict our current data, and they support the proposed direct relationship between the amplitude of the 275-nm CD band and the proportion of BII conformers in DNA. The purported role of exocyclic amines was to increase the number of positive charges near DNA, thus mimicking the direct effect of high salt on DNA bases (58). However, the crosslinking occurred in the DNA major groove near the phosphates and could directly influence the BI/BII balance, which would then affect the CD signal. Electrons in stacked bases may be affected by the BI/BII dynamics stronger than by the electrostatic field of the amines, because the phosphate groups are closer and not shielded from the bases by intervening solvent. This interpretation may explain the effect on the CD of modified DNA caused by sonication (58), which inflicts multiple types of DNA damage and, therefore, may hinder the BI/BII dynamics.

It is possible that the BI-to-BII transitions *per se*, rather than changes in base-pair orientations, are responsible for the steady decrease of the 275-nm CD band (59). The BI-to-BII transitions involve rotations of charged groups (phosphates) along a three-dimensional spiral trace around the base-pair stack, which should affect several electron transitions of the 275 nm band, by modulating the chiral internal electrostatic field. This mechanism was overlooked earlier, because only the BI conformers were originally used for building models of the DNA double helix, and only a handful BII instances was found in refined X-ray structures. In general, the BII conformations are associated with DNA flexibility and high helical twist (60). For natural sequences, the proportion of the BII conformers is zero in A-DNA, reaches nearly 50% in C-DNA, and varies between these limits in B-DNA (5). The driving role of the BI-to-BII transition in defining the intensity of the 275-nm CD band can account for additional puzzling results, such as the contrasting behavior of the 275-nm band versus other CD bands at shorter wavelengths, sequence effects, and other data (27, 61).

The BI-to-BII transitions can occur without changes in the global three-dimensional structure, since the induced increase of base-pair twisting can be compensated by a similar decrease at an adjacent base-pair step. This property allows the rotation of a single base-pair in a stack and takes place even in DNA crystals (62). The increase of positive CD intensity observed below 260 nm (58) can be due to such a compensatory effect in modified DNA. In air-dried DNA fibers, persistent BII conformers should block B-to-A transitions, which explains a striking observation of C-DNA diffraction pattern for NaDNA fibers of this modified DNA under conditions where only A-form is normally observed (23). When the average helical twist is increased by hydration forces or external torques, a higher proportion of BII conformers can serve as an indicator of global structural changes, but this does not mean that the same global changes can be induced by chemical modifications that stabilize BII conformers. Other relevant studies are discussed in the Supplementary Information section.

### C-DNA should be stable in concentrated aqueous solutions of LiCl above 8 M

Finally, we can revisit the question concerning the structure of “a minor variant of the B-form” (36), which gave rise to the original CD spectrum attributed to C-DNA (16). That spectrum was obtained for an unoriented DNA film at 75% air humidity (*a*_w_≈0.75) (16). With respect to our current results (Fig. 1E), it is bound by the spectra of P_L_ in 4.0 and 6.0 M LiCl (corresponding to *a*_w_ of ≈0.79 and ≈0.61, respectively, ref. (35)). The value *a*_w_≈0.75 is reached in 4.5 M LiCl, and the corresponding interpolated spectrum of P_L_ yields a reasonable match to the original reference spectrum (16) (Fig. S3). This result supports the early view that, at high salt concentrations, DNA secondary structure and the associated CD spectra strongly depend on water activity as well as on the direct DNA-ion interactions (20, 63, 64).

In air-dried fibers, C-DNA conformations appear at humidity of 75-79%, and these conformations are characterized by the helical twist of 37.1-37.5°, only 1.1-1.5° higher than in canonical B-DNA (2, 36). As shown here, the transition from a typical B-DNA structure to that found at *a*_w_≈0.75 can be divided into a series of steps, with structural changes similar to those associated with changes in superhelical density (Fig. 3D). Over the entire course of this transition, the helical twist increases by 1.2° (Fig. 3D). This result suggests that the original reference spectrum of C-DNA (16) corresponds to an intermediate on the B-to-C transition pathway rather than “the true C-form”. This intermediate is found at the upper limit of *a*_w_ at which C-DNA may exist. On the other hand, the lower limit of *a*_w_ at which B-DNA diffraction patterns of LiDNA disappear is found at air humidity of 66% (*a*_w_≈0.66) (65). Overall, these results point to a transition zone between *a*_w_≈0.75 and *a*_w_≈0.66, and suggest that the “true C-form” should be found at lower *a*_w_ values.

It is important to note that the environment of DNA in solution is physically very different from that in fibers, as the fiber packing favors extended, approximately straight DNA conformations that have negligible entropy in solution. These geometrical restrictions may favor a higher twist in the B-form and thus facilitate the B-to-C transition (66, 67). If so, the onset of the B-to-C transitions in solution should be shifted from the transition zone in fibers (with *a*_w_ of 0.66-0.75) to a lower *a*_w_. In principle, in saturated solution of LiCl, *a*_w_ is reduced to 0.11 (35), which is substantially lower than *a*_w_ in air-dried fibers featuring canonical C-DNA. Thus, a cognate B-to-C transition should be achievable in concentrated aqueous solutions of LiCl above 8 M. This process should be associated with a further increase in the helical twist, and, as the result, a further decrease of the 275-nm CD band.

## DATA AVAILABILITY STATEMENT

The data underlying this article are available in the article and in its online supplementary material.

## AUTHOR CONTRIBUTIONS

A.K.M. & E.G. conceived and designed the study; A.K.M developed the theoretical framework; M.M. & E.G. performed experiments; A.K.M. & E.G. analyzed the data and wrote the manuscript.

## ACKNOWLEDGMENTS

We thank Dr. Alexander Vologodskii for critical comments, useful discussion, and advice. We also thank Sébastien Brûlé and other members of the Molecular Biophysics facility (Institut Pasteur) for advice and access to the circular dichroism, as well as Sara Chehboub for technical assistance. This work was supported by Agence Nationale de la Recherche (ANR-11-LABX-0011, ANR-11-EQPX-0008, ANR-10-LABX-0062, ANR-19-CE12-0002), Centre National de la Recherche Scientifique (CNRS) and Institut Pasteur.

## SUPPLEMENTARY INFORMATION

### (SI)#SI DISCUSSION

#### Structural interpretation of CD spectra

In supercoiled DNA, short segments involved in plectonemes or other global folds can also deform locally in response to the torsional stress evenly distributed along the double helix, and they should resemble strained segments of deformed minicircles, seen by cryo-EM (34, 68), rather than smooth cylinders. In a double helix with a forced twisted orientation of the opposite ends, the nature of internal deformations is determined by the balance between bending and torsional rigidity. When torsional rigidity is relatively small and bending rigidity is high, the double helix is twisted uniformly, without changing directions of the helical axis. In contrast, torsionally stiff but bendable DNA does not change the twist but adopts the shape of a stretched solenoid or a superhelix, that is, it undergoes solenoidal writhing. In thermodynamic equilibrium, both deformation modes co-exist, and the balance between them shifts from bending to twisting with reduced segment lengths. This property is used in coarse-grained models to separate and quantify the relative weights of *ΔTw* and Wr in Eq. (2). *Wr* is the key parameter in this equation because it can be computed from the three-dimensional coordinates of the closed curve of the helical axis, while *ΔTw* is obtained by subtracting *Wr* from *ΔLk*.

The approach described above may underestimate and neglect the solenoidal writhe at small length scales, meaning that a part of *Wr* is counted as *Tw*. For instance, obtaining an accurate three-dimensional helical trace of long DNAs is difficult experimentally, and so *Wr* is usually computed from the number and geometry of chain crossings in EM images of plectonemes, effectively counting all non-plectonemic writhing as twisting (34, 48, 53). The coarse-grained model in MC simulations calculates *Wr* analytically from the helical axes and, by definition, accounts for all types writhing (49). However, in these simulations, the axis is approximated by a chain of short twistable sticks, with the length depending on the assumed DNA torsional rigidity, which is not precisely known. The reported upper limit of 0.8 for the ratio *qWr*=*Wr/ΔLk* was obtained for the torsional rigidity of 300 pN·nm^2^, and a 30% decrease of this value reduced *qWr* by 10% (49). We are not aware of any studies using larger torsional rigidity, but a 30% increase likely gives a similar effect of opposite sign and *qTw* < 0.1, which would be not far from the value for 0.5 M LiCl in Fig. 2C. The uncertainty of ±30% is smaller than the range of the experimental values of torsional rigidity previously obtained by different methods (69).

#### Contrasting views on the structure of DNA in concentrated ionic solutions

The earlier CD-based results reporting the effects of common salts on the helical twist were verified by measuring twist with alternative methods. Anderson and Bauer (51) added LiCl (up to 300 mM) to topoisomerase relaxation reactions and obtained distributions of circular DNA topoisomers that had positive *Δ Lk* values in a gel electrophoresis buffer used for the analysis of topoisomers by band counting. Their results largely agreed with the CD data, suggesting that the intensity of the 275-nm band reflected changes in twist (31). However, it was also reported that, at salt concentrations above 300 mM, the helical twist actually ceased to increase and, perhaps, even had a tendency to decrease (51). More recently, the same set of salts and concentrations were tested directly using single-molecule magnetic tweezers (52). In low-salt conditions, changes in twist were similar to the earlier data (51), but showing no signs of saturation over the range of LiCl concentrations from 50 mM to 1 M. Thus, the earlier reported saturation effect (51) could have resulted from incomplete relaxation of supercoiling due to the inhibition of topoisomerase activity by relatively high salt.

The effect of salt concentrations above 3 M was tested using the ethidium titration method, which measures the number of the bound ethidium molecules required to unfold a negatively supercoiled DNA to a circle with *Δ Lk*=0 (70, 71). Wang and colleagues were the first to apply this method to measuring the helical twist of DNA in 3 M CsCl (72). Baase and Johnson (22) used concentrations above 3 M and found almost identical twist values for 6.2 M LiCl and 3 M CsCl, suggesting that the twist did not increase, if not decreased, between 3 M and 6.2M LiCl. However, this study did not take into account earlier studies reporting a salt-dependent transition between two conformational states of supercoiled DNA (54, 55, 71, 73, 74), later rediscovered as loosely and tightly interwound superhelices (34, 53). In ethidium titration experiments, this transition is accompanied by a change in sedimentation rates, which can be confused with the principal minimum at *Δ Lk*=0 as previously reported by others (55, 73). One cannot exclude that in 6.2 M LiCl, the ethidium concentration was erroneously determined for a transition between the two states of supercoiled DNA (loosely versus tightly interwound superhelices) and not for unfolding to *Δ Lk* =0. This alternative conclusion, in particular, is justified by the apparent absence of dependence of minimal sedimentation rates on the number of bound ethidium molecules. For flat circles, these rates should decrease due to increasing DNA length and other unavoidable physical factors (55), as had been shown in previous studies and also by the same authors in the verification experiments with 3 M CsCl (22). Sedimentation data including nicked circles in the entire range of salt concentrations could have provided a reasonable control experiment, but they were not available (22).

## SI FIGURE LEGENDS

**Figure S1.**
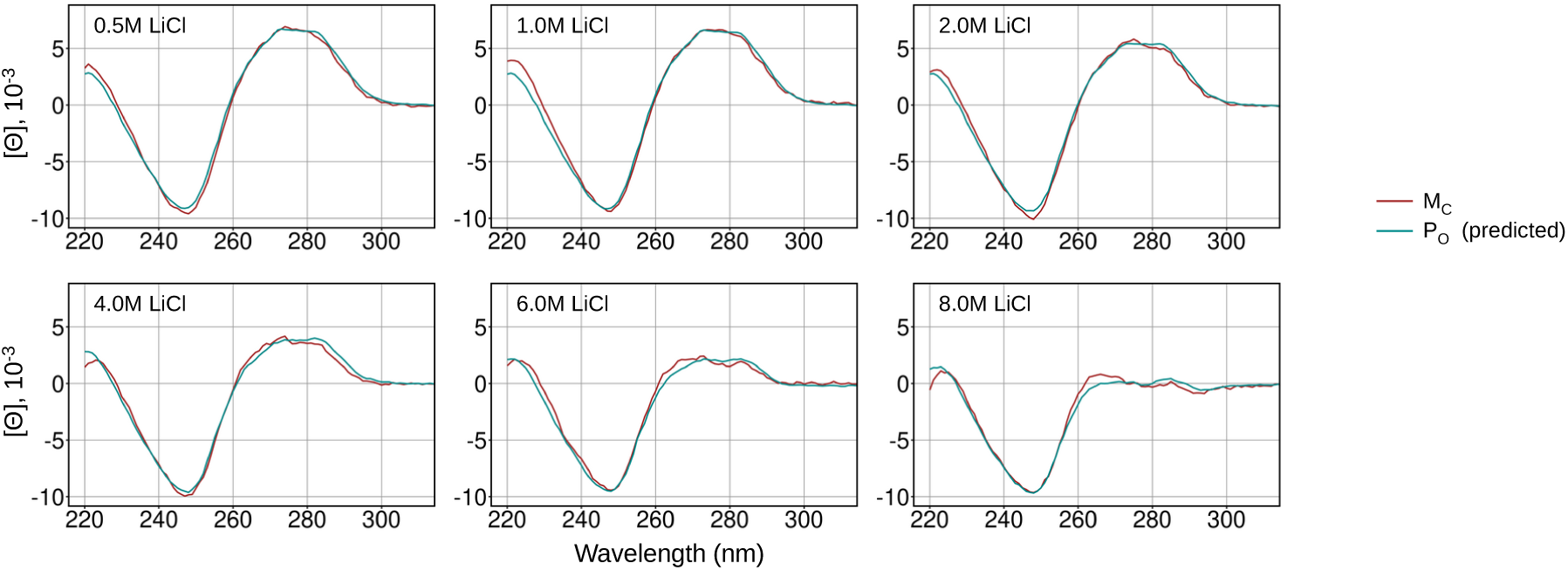
CD spectra of M_C_ versus predicted CD spectra of P_O_. P_O_ spectra were computed by linear interpolation from the neighboring experimental spectra to LiCl concentrations of optimal fit (as in Fig. 3A). The latter were predicted by minimization of the root-mean square difference between the computed P_O_ spectra and target M_C_ spectra. The concentrations of LiCl for M_C_ spectra are indicated on each plot. The corresponding concentrations of LiCl for the predicted P_O_ spectra are 0.301 M, 0.334 M, 0.763 M, 1.532 M, 2.972 M and 4.894 M, respectively.

**Figure S2.**
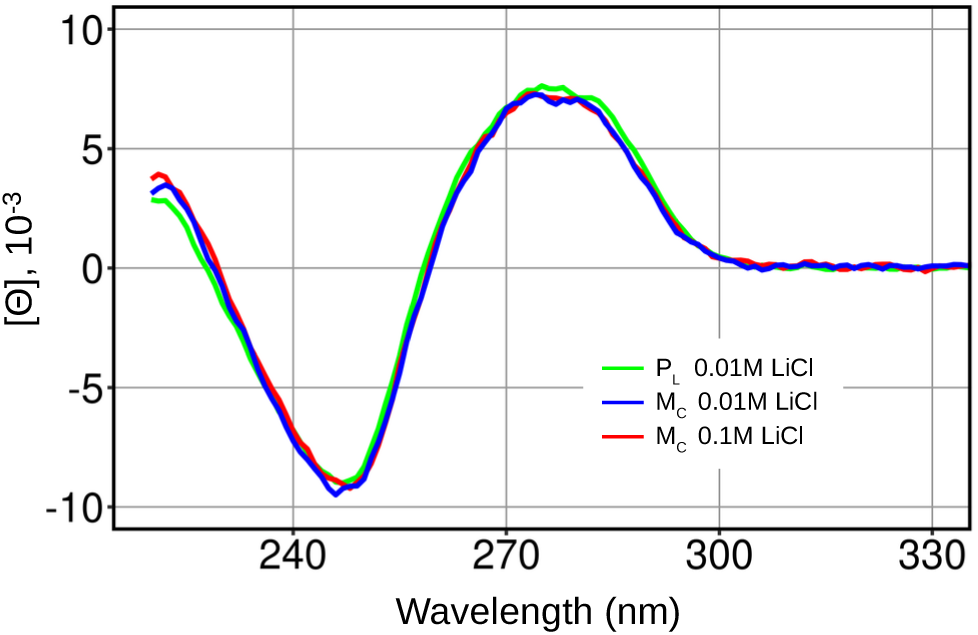
CD spectra of M_C_ and P_L_ at low LiCl concentrations. Concentrations of LiCl are indicated. P_L_ data were re-plotted from Figure 1E.

**Figure S3.**
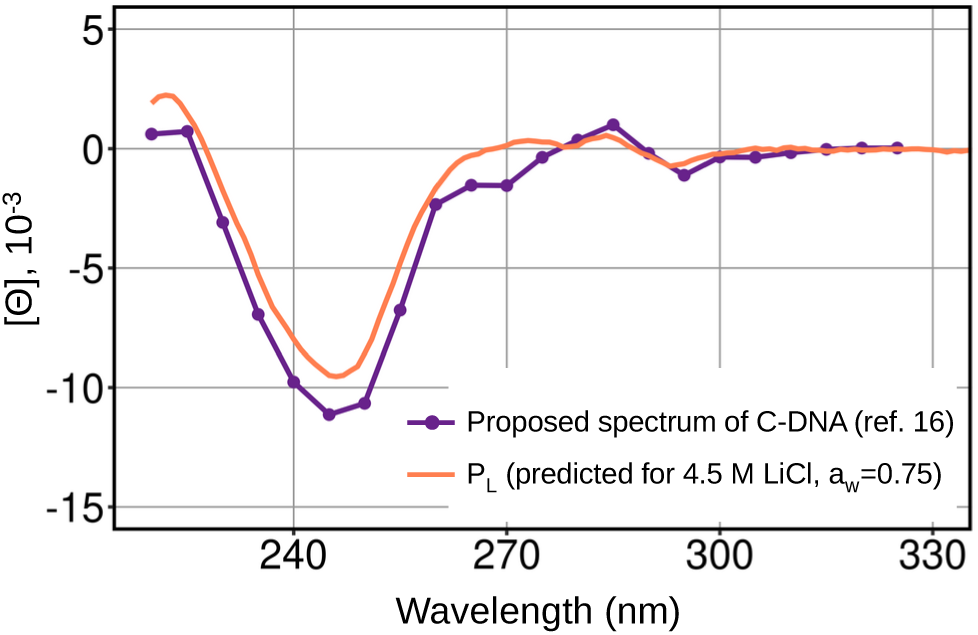
The reference spectrum for C-DNA versus the predicted spectrum of P_L_ at a_w_=0.75. The CD spectrum of an unoriented DNA film at 75% air humidity, previously proposed as a C-DNA standard (16, 35), is compared with the spectrum of P_L_ predicted for a solution of LiCl with a_w_=0.75 using linear interpolation from the appropriate experimental spectra (as in Fig. 3A).

